# Amino acid-derived quorum sensing molecule alanine on the GIT tolerance of the lactobacillus strains in the co-cultured fermentation model

**DOI:** 10.1101/2021.06.01.446689

**Authors:** Jianzhu Wen, Lei Cui, Mengqi Chen, Qiang Xia, Xiaoqun Zeng, Yuxing Guo, Daodong Pan, Zhen Wu

## Abstract

As the importance of gut microbiota in health is increasingly recognized, the interest in interventions that can modulate the microbiota and its interactions with its host has soared. The survival status of the probiotics in the gastrointestinal environment and the microbial interactions between the LAB have also received considerable attention. In the present research, the gastrointestinal environment tolerance, adhesion ability, and biofilm formation of the lactobacillus strains in the co-culture system were explored, through the real-time fluorescence-based quantitative PCR, UPLC-MS/MS metabolic profiling analysis and Live/Dead^®^ BacLight™ cell staining methods. The results show that the co-culture system can promote the release of signal molecules and can effectively protect the liability of the *Lactobacillus acidophilus* in the gastrointestinal environment. Meanwhile, amino acid-derived quorum sensing molecule L-alanine (1 %) can effectively enhance the communication of the cells in the complex fermentation model, which leads to the increase of the liability of the *L. acidophilus* in the gastrointestinal environment.

**Importance:** The findings of this study provide a clue to the amino acid-derived metabolites in the communication among cells in the GIT environment, which can enhance the communication of the lactobacillus strains in the complex fermentation model. Meanwhile, the liability of *Lactobacillus acidophilus* could be enhanced in the co-culture system during the gastrointestinal environment stress with the amino acid-derived quorum sensing (QS) components, and it will shed some light to the application of the amino acid-derived QS molecules in the fermentation stater industry.

## Introduction

Probiotics are defined as living microorganisms when given in sufficient amounts that can improve the balance of microorganisms [1], especially in the gastrointestinal tract. Producing beneficial effects on the health of the host [2]. Most of the probiotics consist of *Saccharomyces boulardii* yeast or lactic acid bacteria (LAB), and the widely used *Lactobacillus* and *Bifidobacterium* [3]. It mainly exerts its beneficial effects by reducing the invasion of pathogenic microorganisms and changing the host immune response in the gastrointestinal tract (GIT).

LAB is a universal physiological flora that exists in the human intestinal tract [4]. It can colonize in the human intestine and form a protective film to prevent the impact of the harsh gastrointestinal tract (GIT) micro-environment on the bacteria [5]. Meanwhile, the short-chain fatty acids and lactate produced by LAB have been shown to play an important role in maintaining intestinal homeostasis [6]. In addition, LAB can improve the nutritional value of the fermented foods, decrease intestinal infections and serum cholesterol levels [7]. Compared with other probiotics, LAB is more resistant to acids and bile salts in the GIT environment [8]. For this purpose, LAB including *Bifidobacterium*, *Lactobacillus* are more popular in the fermentation food industry [9], and the models of co-culture fermentation with some outstanding probiotics are more common in the current fermented dairy products. However, there are still a lot of questions exist of how the different strains interact with each other in this special matrix.

It has long been appreciated that there is a special mode of interaction in the bacterial groups through the exchange of extracellular signaling molecules in quorum sensing [10, 11] and there is a regulatory mechanism in this communication process between bacterial groups and many important physiological functions such as bioluminescence, motility, production of secondary metabolites and biofilm formation [12–14]. Bacterial type, bacterial density and environmental stimuli, including acid-base stress, bacterial surface adhesion have been reported to affect the expression of QS-related signaling molecule [15]. In Gram-negative bacteria, N-acyl homoserine lactone is the most common signaling molecule [16]. While studies in recent years have found that the most common signaling molecules in Gram-positive bacteria are signaling peptides (AIP) and autoinducer-2 (AI-2) [17–19]. AIP is transduced by a two-component condition of transmembrane histidine kinase (HK) and transcription regulator (RR) [14]. AgrD encodes the precursor of the AIP, whereas the integral membrane protein AgrB is involved in its processing and secretion of the thiolactone-modified cyclic oligopeptide [20]. AI-2, a novel furanosyl borate diester, is amplified by the *luxS* (S-ribosylhomocysteine cleavage enzyme) gene product in the genome of many gram-negative and gram-positive bacteria. This has been proposed as a ‘universal’ signaling molecule for interspecies communication [12, 21]. In the metabolic process, the use of SAM as a methyl donor produces the intermediate S-adenosylhomocysteine (SAH), which is hydrolyzed to S-ribosylhomocysteine (SRH) and adenine by the nucleosidase enzyme Pfs (5’methylthioadenosine/ S-adenosylhomocysteine nucleosidase). LuxS catalyzes the cleavage of SRH to 4, 5-dihydroxy 2, 3-pentanedione (DPD) and homocysteine [22].

In recent years, QS related cell-cell communication in lactic acid bacteria has received more and more attention, especially in the co-culture condition. As we all know, the main metabolites of lactic acid bacteria are amino acids, extracellular polysaccharides and lactic acid. Some research also found that quorum sensing affects the type and abundance of bacterial metabolites in the co-culture condition. Meanwhile, whether the type and abundance of bacterial metabolites will also affect the synthesis of AIP and AI-2 is still worth further studying.

This study explored the QS network of the lactobacillus in the co-culture condition, and the biofilm changes, metabolite profiles and the liability of the lactobacillus strains in the simulated gastrointestinal conditions were also investigated according to the amino metabolites-mediated QS changes. It is hoped that this study can further the understanding of the regulatory mechanism of quorum sensing on the growth and behavior of the strains in the gastrointestinal micro-environment. And to provide insight to whether there is a link existing between the intestinal tolerance of LAB and the influence of certain amino acids in the synthesis of AI-2 in the QS system.

## Materials and methods

### Strains and culture conditions

*Lactobacillus acidophilus* CICC6074 was obtained from the China General Microbiological Culture Collection Center (Beijing, PR China). *Sterptococcus thermophilus* ABT-T and *Lactobacillus delbrueckii* subsp. *bulgaricus* BNCC336436 were stored in our laboratory. For the experiment, all the cells are cultured in MRS media at 37 °C for 12 h before use.

### Growth characteristics of the strains in co-culture condition

Exploring the effects of the strains under mixed-culture condition, *L. acidophilus* and *L. bulgaricus* and *S. thermophilus* were cultured in 37 °C MRS medium under the same conditions with different inoculation ratios. The ratio of *L. acidophilus* and *L. bulgaricus* and *S. thermophilus* were 1:1:0, 1:1:1, 1:1:2, 1:1:3. The growth of the strains under the mixed strain culture conditions was measured at 600 nm every 2 hours, and the pH value of the acid production was measured with a digital pH meter within the 24 hours’ culture time.

### Relative account of the different strains in the co-culture condition

Quantitative analysis of account of the *L. bulgaricus*, *S. thermophilus* and *L. acidophilus* in the fermentation broth at different fermentation time in the co-culture condition by real-time PCR with the gene-specific primer design by Primer Premier 5.0 software.

### AI-2 bioassay detection

Autoinduction of luminescence in the marine bacteria *V. fischeri* and *V. harveyi* was described in the early 1970s [23]. In this study, the AI-2 reporting strain *V. harveyi* BB170 was adopted for the signaling molecules detection and the AI-2 bioassay was performed according to the method as previously described [9]. Cell free culture fluid (CF) was prepared as follows: after centrifuging the test strain at 4600 g for 10 min, and the resulting supernatant was further filtered through a 0.22 μm filter to obtain the CF sample. The reporter strain *V. harveyi* BB170 was diluted 1: 5000 in fresh AB medium, and then 99 mL of AB culture was mixed with 1 mL of the CF sample fluid and incubated at 30 °C. The fluorescence intensity was detected every 1 hour with the black 96-well microplate reader. Experiments were performed in triplicate and repeated at least three times.

### Quantitative reverse transcription PCR for the detection of the *LuxS* signaling molecule gene

The strains *L. acidophilus*, *L. bulgaricus* and *S. thermophilus* (1:1:1) were inoculate in the MRS medium at 37 °C for 18h. Then, the cells were collected for the RNA extraction using Magen kit (Magen Biotech Co.,Ltd., Guangzhou, China). After that, the extracted RNA was reverse transcribed with the kit (TranScript All-in One First-Strand cDNA Synthesis SuperMix for qPCR, TransGen Biotech, Beijing, China) for real-time PCR analysis with the internal reference gene is *dp3d* (DNA polymerase Ⅲ, delta subunit).

Gene-specific primers F (5’-GAATGTGGGCGTTAAGCAAACC-3’), R (5’-TGCACGTTCCTCATCACTATCG-3’). Use the purified cDNA and gene-specific primers for real-time PCR analysis in triplicate with the kit (TransStart Tip Green qPCR SuperMix, TransGen Biotech, Beijing, China). Finally, the number of cDNA was normalized to the abundance of 16S rRNA by the 2^−ΔΔCT^ method and the relative quantitative method was used for *LuxS* signaling molecule gene analysis.

### Metabolomics analysis

The harvested cocultured LAB (18 h) were centrifuged (4600 g, 5 min) and washed twice with double distilled water. Then the supernatant was removed (4600 g, 5 min) and the bacterial samples were lyophilized by vacuum-freeze drying. Bacteria samples (10 mg) were resuspended in 1.5 mL pre-cooled methanol and subjected to three freeze-thaw cycles before sonication in an ice bath for 15 min (cycles: 1 min pulse followed by 1 min pause). Aliquot (10 μL) was vortex-mixed with 10 μL of NEM solution (20 mM) in phosphate buffer (0.1 M, pH7.0) containing 10 mM ascorbic acid, 10 mM EDTA and 7% DMSO for 1 min. 10 μL tBBT solution (0.23M in DMSO) was added followed with the addition of 87.5 μL borate buffer (0.2 M, pH 8.8) containing 20 mM TCEP and 1 mM ascorbic acid. After vortex-mixing and standing for 2 min, 33 μL 5-AIQC solution was then added and incubated at 55 °C for 10 min. The mixture was cooled down to the ambient temperature and added with 2 μL formic acid followed with solution was filtered by a 0.22 μm membrane filter before UPLC-MS/MS analysis.

The UPLC-MS/MS consisted of an Agilent 1290 UPLC coupled to an Agilent 6470 triple quadrupole mass spectrometer equipped with an electrospray ionization (ESI) source (Agilent Technologies, USA). The 5-AIQC-tagged samples (1 μL) were individually injected on an UPLC column (Agilent ZORBAX RRHD Eclipse XDB C18 column, 2.1 × 100 mm, 1.8 μm particles) with its temperature set to 50 °C. Water and methanol containing 0.1% (v/v) formic acid were used as two mobile phases A and B, respectively, with flow rate of 0.5 mL/min. An optimized gradient elution scheme was employed as 1% B (0-2 min), 1-3.8% B (2-4 min), 3.8-14% B (4-7.3 min), 14-22% B (7.3-10.7min), 22-24% B (10.7-14.7 min), 24-30% B (14.7-16 min), 30-60% B (16-16.3 min), 60-70% B (16.3-17.3 min), 70-95% B (17.3-17.31 min), and 95% B (17.31-20 min). Electrospray ionization was performed in the positive ion mode using N^2^ at a pressure of 50 psi for the nebulizer with a flow of 10 L/min and a temperature of 315 °C, respectively. The sheath gas temperature was 350 °C with a flow rate of 10 L/min. The capillary was set at 4000 V. Multiple reaction monitoring (MRM) has been used for quantification of screening fragment ions.

The metabolism experiments were repeated six times and the quality control samples (QC samples) were analyzed in continuous random order. Peak determination and peak area integration were performed with MassHunter Workstation software (Agilent, Version B.08.00). Standard curves were constructed by least-squares linera regression analysis using the peak area ratio of derivatized individual standard against the nominal concentration of the calibrator. Quantification of samples was performed identically. The OriginPro 9.1 and SIMCA 14.1 was used for multivariate statistical calculations and plotting.

### Survival rate of the strains under simulated gastrointestinal fluid

Simulated gastrointestinal digestion according to the method of Ranadheera (2014) with some modifications. Bacteria (*L. bulgaricus*, *S. thermophilus* and *L. acidophilus*) cultured in 37 °C MRS medium for 18h at different inoculation ratio were washed three-times, and then treated with artificial gastric fluid for 2 h and simulated intestinal fluid for 3 h at 37 °C. Serial dilutions of suspended bacteria were plated on MRS agar. Survival rates were calculated with three independent experiments.

### Aggregation Assay

The aggregation assay was according to the method of Cao (2019) with some modifications. Different experimental groups (1:1:0 and 1:1:1) were grown in MRS medium for 18 h at 37°C, centrifuged at 4600 g for 10 minutes, washed twice with PBS buffer. Then adjust the suspension to an optical density value of 1±0.05 at 600 nm (A_0_). The bacterial suspension (4 mL) mix was incubation at room temperature for 4h, and then the absorbance was measured at 600 nm (A_4_). The self-aggregation rate is calculated as follows:

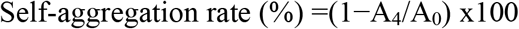

where A4 and A0 represent the absorbance at 4h and 0h, respectively. The aggregation assay of the bacteria with the gastrointestinal fluid treatment was calculated with the same experimental procedure above.

### Adhesion assay *in vitro*

The adhesion assay was carried out with the Caco-2 cell model. After the gastrointestinal treatment, the LAB samples were stained with the 6-carboxyfluorescein diacetate (CFDA, 10 μM, Sigma, Darmstadt, Germany). When the Caco-2 cells covered 80% of the area of the petri dish bottom, the stained bacteria were co-cultured with the Caco-2 cells for 2h at 37°C. Finally, fluorescence intensity (excitation wavelength 485 nm; emission wavelength 538 nm) was measured using Tecan Infinite M200 Pro (Shanghai, China) and the adhesion rate was expressed as the percentage of fluorescence after binding to Caco-2 cells relative to the bacterial suspension added to the wells.

### Biofilm formation detection

The biofilm formation of the of the strain in the co-culture condition was detection according to the method of Rieu (2007). The expanded *L. bulgaricus*, *S. thermophilus*, and *L. acidophilus* (1×10^8^ CFU/mL) were added to a 24-well plate at different ratios and cultured at 30°C for 24 h. Bacteria that not bound to the well plate was washed with 150 mM NaCl solution. For the stimulated gastrointestinal environment analysis, the same amount of the gastric juice or intestinal juice were added to the well plate and incubate for 2 h, and then wash twice with NaCl solution (150mM). For the staining procedure, the control group and the treatment group were stained with crystal violet for 45 min, after washing the unbounded crystal violet, 96% ethanol was added to dissolve the crystal violet for another 45 min, and then the absorbance was measured with a microplate reader at 595 nm.

### Cell viability test

The Live/Dead^®^ BacLight™ cell viability test kit can reliably and quantitatively quickly distinguish live bacteria from dead bacteria. Bacteria with intact cell membrane will show green fluorescence, while bacteria with damaged cell membrane will show red fluorescence. The cell viability was detected after the stimulated gastrointestinal condition treatment. Mix equal volumes of Syto 9 and propidium in a centrifuge tube, then 3 μL of dye mixture was added to the bacteria suspension and incubated for 30 min in the dark. The image of the Live/Dead bacteria was observed using a fluorescent inverted microscope (Olympus, Japan).

### Statistics

Data were analyzed using the one-way analysis of variance (ANOVA) and independent-samples test by SAS 9.1 software with *P*< 0.05 considered significant. Multi-Experiment Viewer (MeV) software was used to quantification of different LAB in the co-culture condition by the microarray heatmap which combined with hierarchical cluster analysis (HCA). The unsupervised model (PCA) and partial least squares discrimination analysis (PLS-DA) model was established to evaluate the impact of metabolites by using SIMCA-P software.

## Results

### Growth characteristics of the strains in co-culture condition

It can be seen from OD_600nm_ that the bacterial density is lower when *S. thermophilus* and *L. acidophilus* are cultured separately while the bacterial density was increased when the bacteria cultured together. However, there is no significant difference in the changes of the mix strain ratio. Besides the OD value comparison, the abundance of different strains was also quantified using real-time fluorescent PCR method. The results showed that the abundance of strains is significantly different with the ratio of the strains. In the mono-bacterial fermentation condition, *L. bulgaricus* has the highest OD value compared with *L. acidophilus* and *S. thermophilus*. Meanwhile, the growth rate of *S. thermophilus* was the lowest one. It shows that in the co-culture condition, both the OD value and pH value were more significant compared with the mono-fermentation samples (Figure 1A and 1B). When the inoculation ratios of *L. bulgaricus*, *S. thermophilus* and *L. acidophilus* are 1:1:1, 1:1:2 and 1:1:3, the abundance of *L. bulgaricus* is decreased in the starters while, the abundance of *L. acidophilus* is significantly increased. It shows that proper increase of *L. acidophilus* can promote the growth of other strains in the starter. Meanwhile, according to the result of the AI-2 assay, the fluorescence value in the mix culture condition is also higher than the mono-culture groups (Figure 1C and 1D).

**Figure 1.**
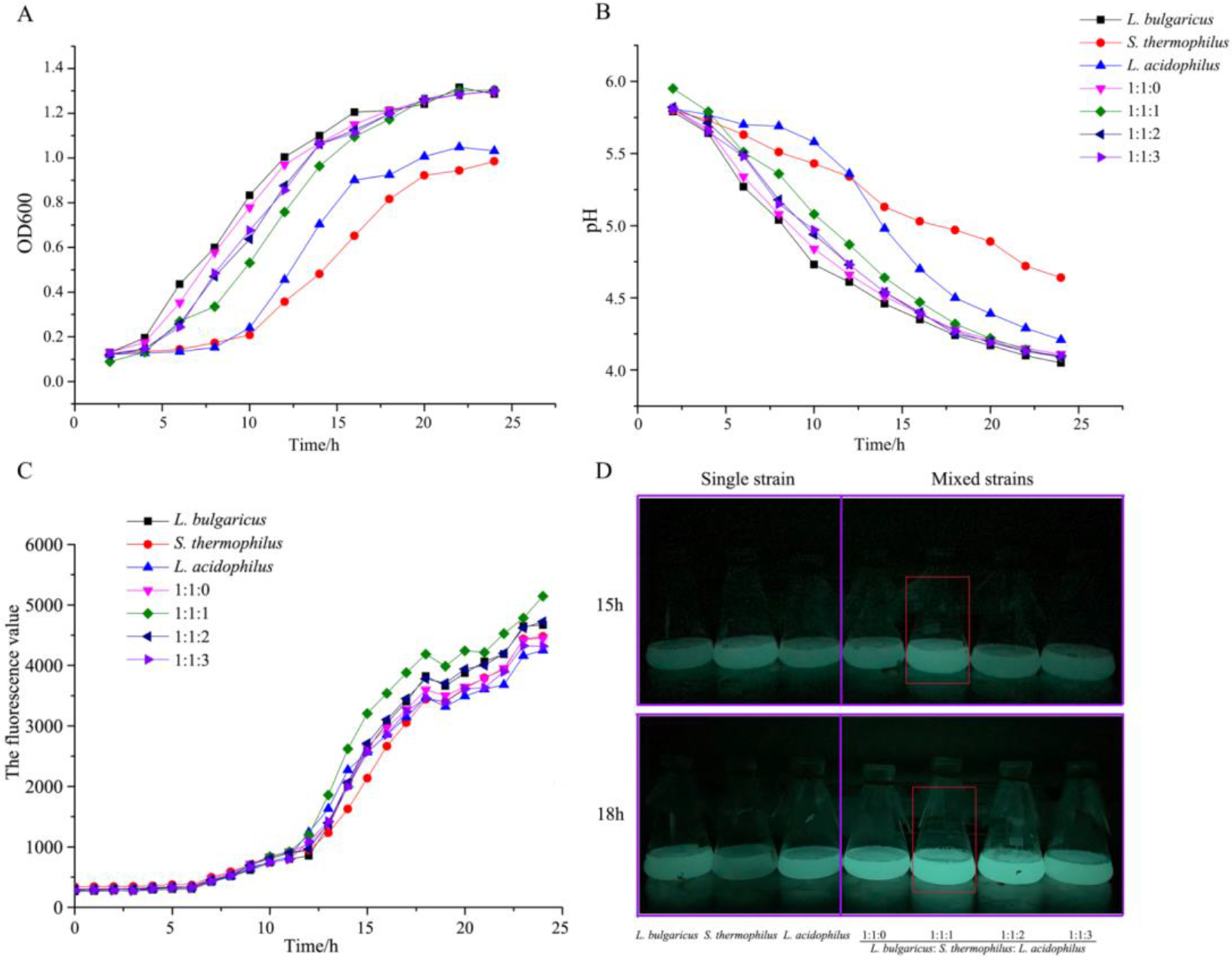
The screening process of the lactobacillus strains in the co-culture condition. A, the OD_600nm_ of co-cultivation strains; B, the pH value of culture media during the culture condition; C, the fluorescence value of AI-2 in the co-cultivation condition; D, the fluorescence image of the culture media in the single and mixed culture condition.

### The tolerance of the LAB strains in the QS related stimulated GIT condition

In the QS related stimulated GIT condition, *L. acidophilus* can increase the resistance of the strain to gastrointestinal fluids. The fluorene density of AI-2 in all the groups were increased with the culture time from 0h to 24h, the AI-2 didn’t change too much after 24h. The fluorene density of 1:1:1 group were highest among other treated groups, which is almost 5500. Meanwhile, compared with the 1:1 group, the fluorene density value was significantly higher in 1:1:1 group that *L. acidophilus* strain presented which means the co-culture condition is good for the release of AI-2 when cultured with *L. acidophilus* (Figure 2A and 2B). This result can also be proved in the Figure 2C, which shows the relative number of the three strains in the co-culture condition. Here the number of *L. acidophilus* is much higher than other two strains. After the gastrointestinal fluid treatment, the liability of the strains in the co-culture condition, especially the 1:1:1 group, were much higher than other groups, though the AI-2 activity between the two treated group was basically the same (Figure 2D). Furthermore, the *LuxS* gene involved in the synthesis of AI-2 was verified by real-time PCR. The results showed that the *LuxS* genes of *L. bulgaricus* and *L. acidophilus* were up-regulated at 18 h compared to 12 hours of cultivation (Figure 2E).

**Figure 2.**
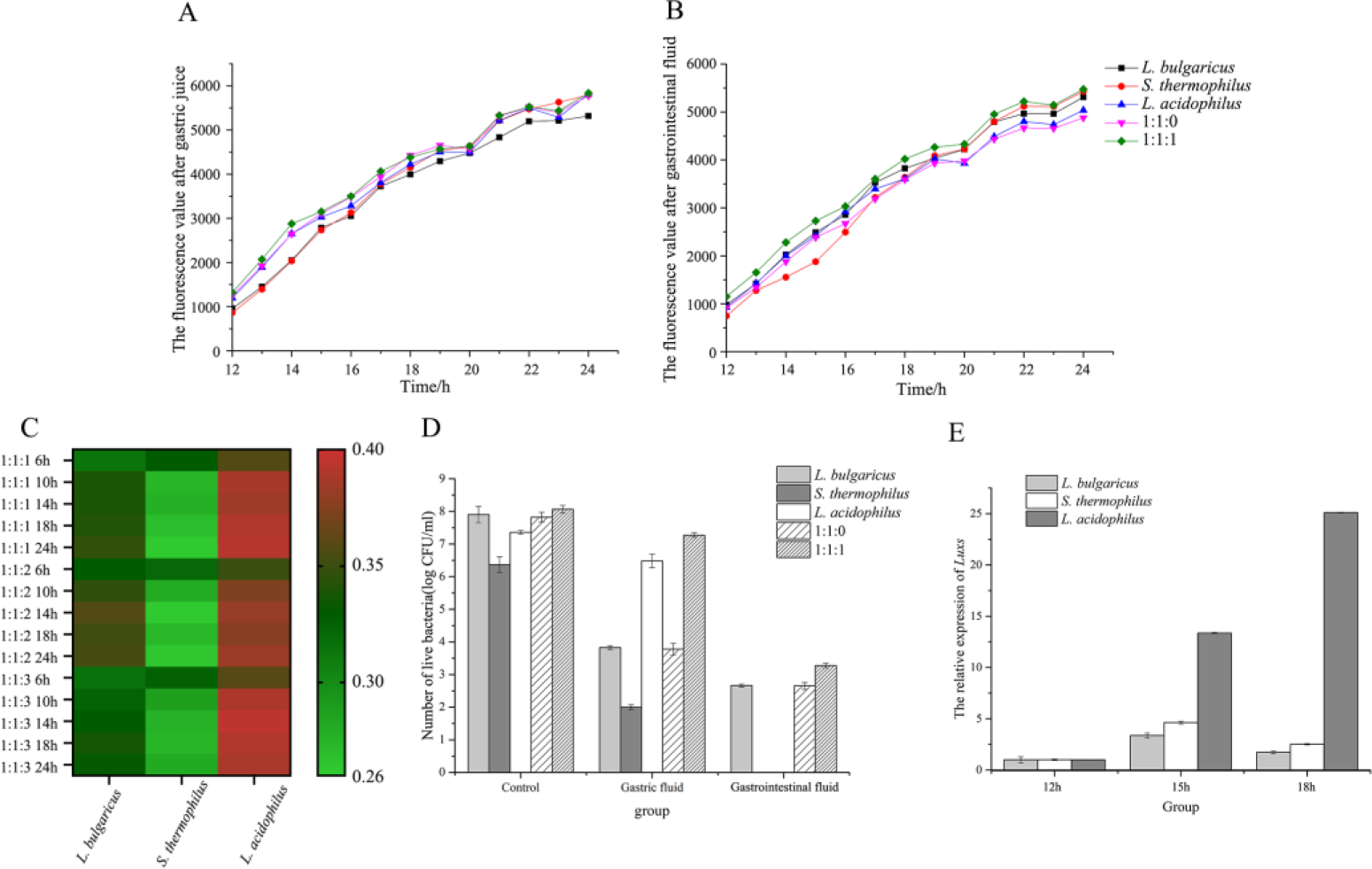
The influence of gastrointestinal condition on the growth characteristics of bacteria under co-culture conditions. A, the fluorescence value of the mixed strains in co-culture condition after gastric juice treatment; B, the fluorescence value of the mixed strains in co-culture condition after gastrointestinal fluid treatment; C, the ratio of the single strain under the co-culture condition; D, the number of live bacteria in the gastrointestinal fluid under co-culture conditions; E, the *LuxS* gene expression of the strains under the co-culture conditions.

### Metabolomics analysis

In the multivariate statistical analysis of the metabolomics profiles, we investigated the changes of amino metabolites of different lactic acid bacteria. The PCA score chart (Figure 3A) shows the obvious metabolite spectrum of *L. acidophilus*, *L. bulgaricus*: *S. thermophilus* 1:1, *L. bulgaricus*: *S. thermophilus*: *L. acidophilus* 1:1:1. The result of PCA score chart can directly explain the metabolic differences of the samples. At the same time, the heat map of the correlation between different metabolites shows whether the relationship between each metabolite is positive or negative (Figure 3B). Amino acid such as L-alanine and L-proline is positive to the SAM and SAH, some are negative such as L-glutamic and L-cystine. In addition, stacked histogram analysis showed the concentration of each significant metabolite by the multivariate statistical analysis (Figure 3C). The metabolites pathway (Figure 3D) related to the producing of AI-2 also given according to the Metabolomics analysis, which L-alanine promotes the synthesis of Homocysteine and further synthesizes SAM and SAH.

**Figure 3.**
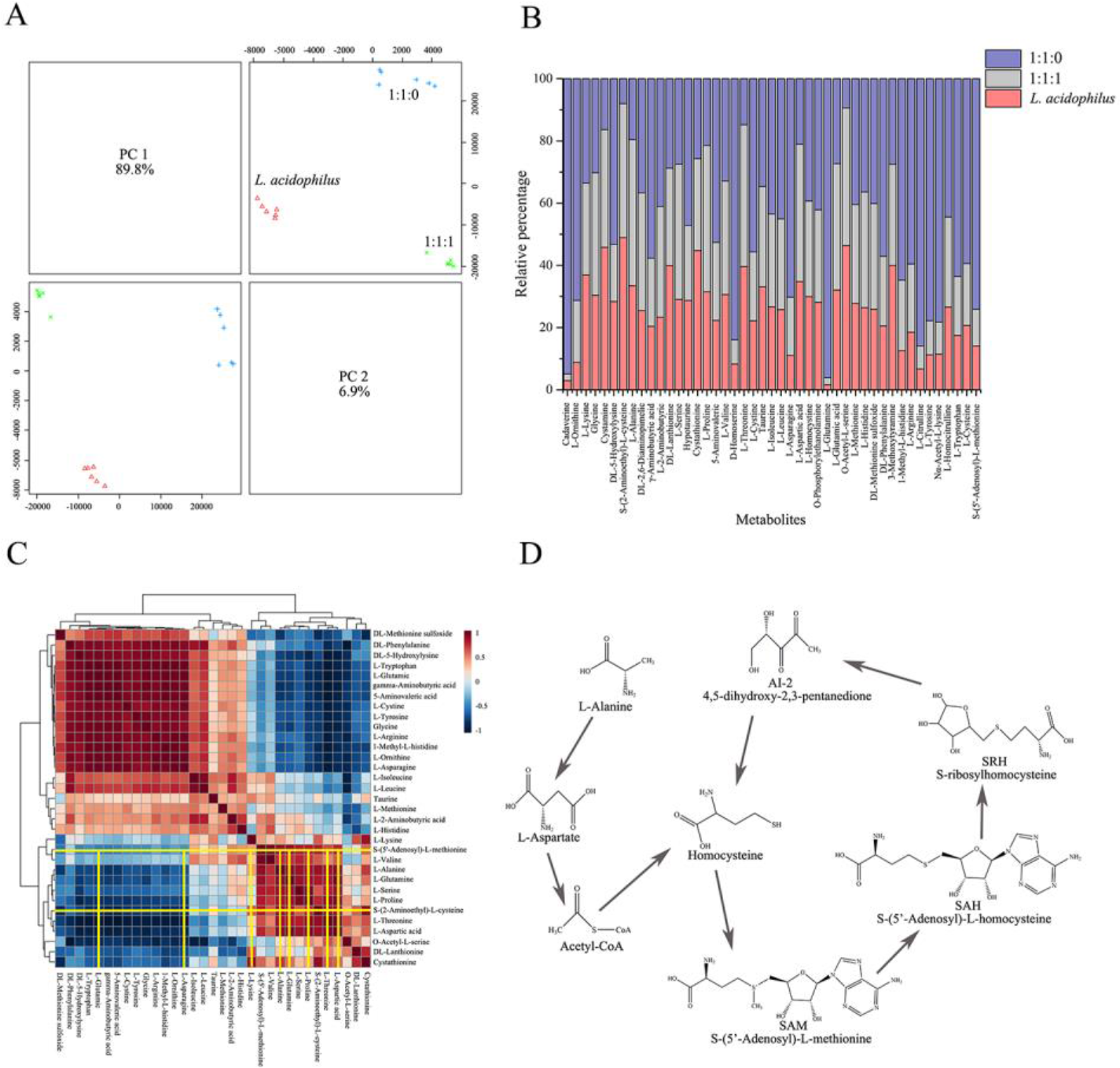
Amino metabolites analysis of culture media under the co-culture conditions. A, the PCA analysis of metabolites in the different mix-culture ratios; B, the concentration of different metabolites in the different mix-culture groups; C, the correlation analysis of the differential metabolites; D, the synthesis pathway of SAM and SAH in the amino acid metabolites pathway.

The variable importance of projection (VIP, VIP obtained by PLS-DA model) (Figure 4A) value combined with the P value was used as the screening criterion to obtain significantly different metabolites. If the metabolite meets the conditions of VIP>1 and P value<0.05, then it is a significantly different metabolite. All these differential metabolites are given in the table S1. Among the eight different metabolites, L-glutamic acid, L-asparagine and L-lysine are not positively correlated with S-(5’-Adenosyl)-L-homocysteine and S-(5’-Adenosyl)-L-methionine. Compared with the concentrations of the other 5 metabolites, L-alanine is a significantly different metabolite. Through the KEGG metabolic pathway, L-alanine was found to play an important role in the synthesis of S-(5’-Adenosyl)-L-homocysteine and S-(5’-Adenosyl)-L-methionine (Figure 3D).

**Figure 4.**
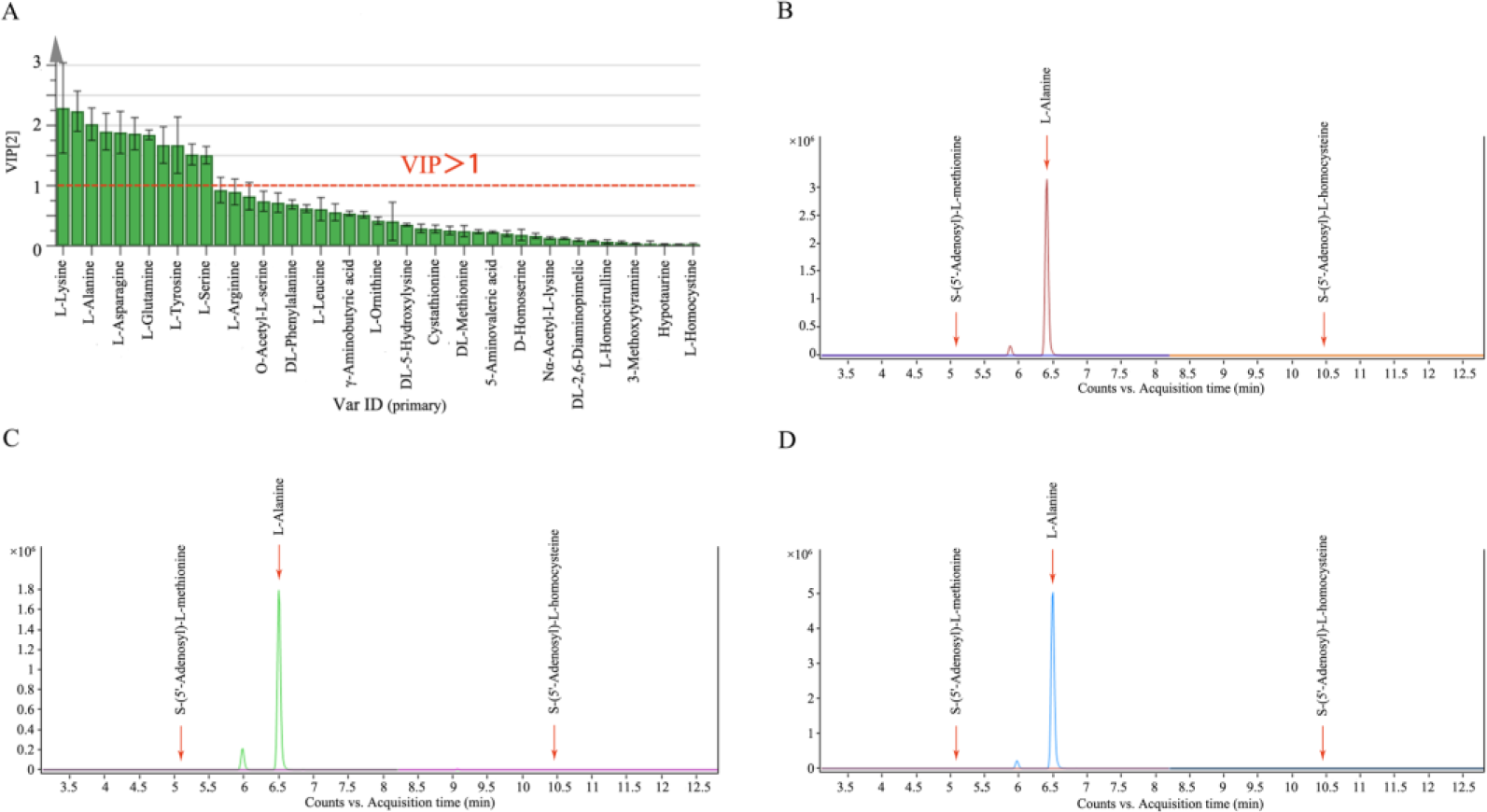
VIP analysis and chromatogram of differential metabolites. A, the VIP analysis of metabolites; B, the chromatograms of SAM, L-alanine and SAH in *L. acidophilus*; C, the chromatograms of SAM, L-alanine and SAH in 1:1:0 fermented group; D, the chromatograms of SAM, L-alanine and SAH in 1:1:1 fermented group.

### Amino acid-derived L-alanine on the AI-2 release and growth characteristics of the strains in the co-culture condition

In this study, strain *V. harveyi* BB170 was used to detect the ability of the co-culture condition to secrete AI-2 molecules when cultured with different concentrations (1%, 2% and 3%) of L-alanine. The results showed that when cultured with 1% concentration of L-alanine, the secretion of AI-2 molecules was significantly increased compared to the one without L-alanine. After 22 hours of culture time, AI-2 molecules activity tends to be stable (Figure 5A). However, the bacterial density at OD600nm did not increase after adding 1% L-alanine (Figure 5B). Similarly, the acid production capacity was not significantly enhanced (Figure 5C). While, the addition of 1% L-alanine can significantly promote the growth of *L. acidophilus* which can be seen from the relative number of the strains in the heatmap illustration (Figure 5D).

**Figure 5.**
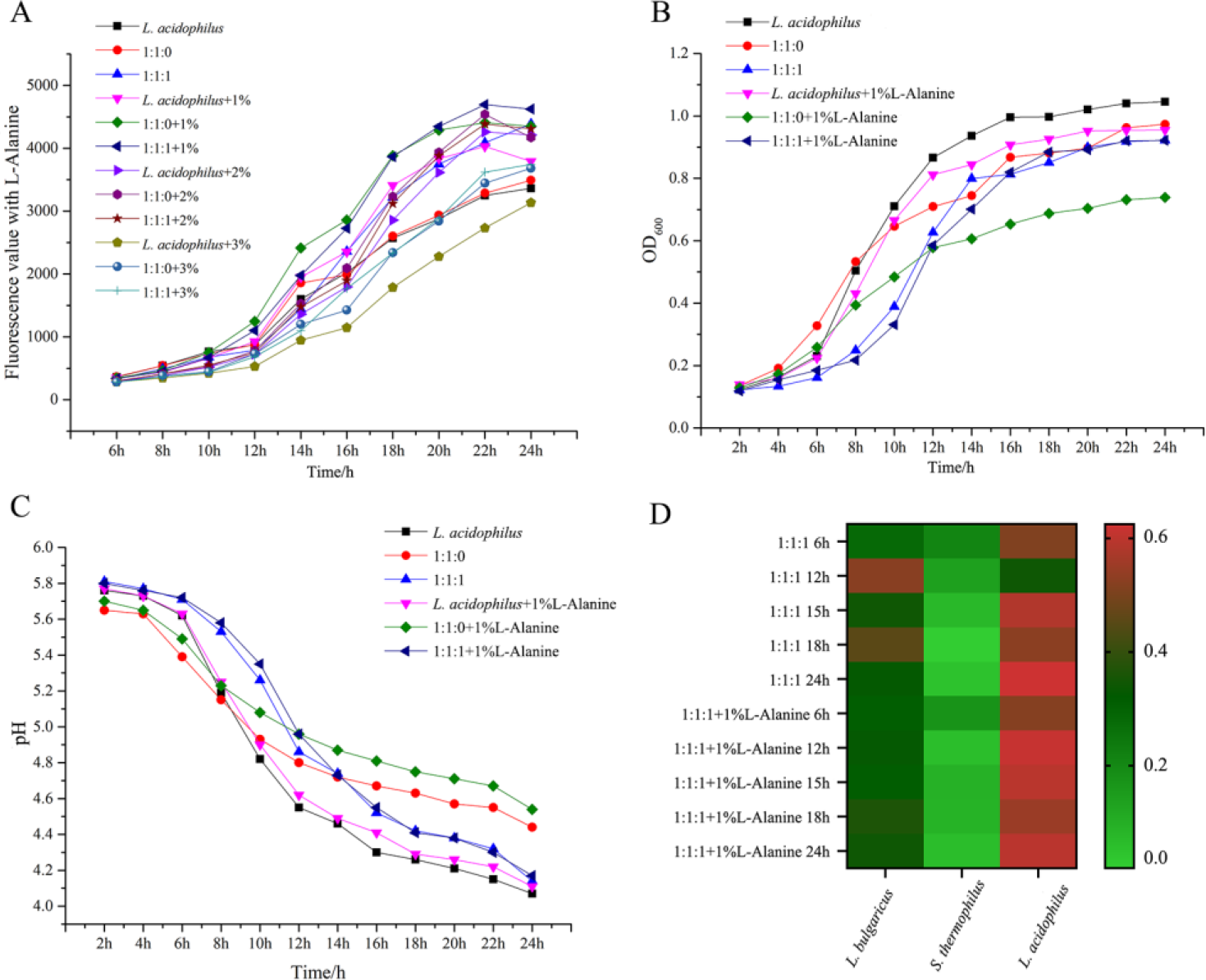
Fluorescence value and bacterial growth of co-culture condition pretreated with L-alanine. A, the Fluorescence value of co-culture media when pretreated with L-alanine; B, the OD_600nm_ of co-culture media when pretreated with L-alanine; C, the pH value of co-culture media when pretreated with L-alanine; D, the bacterial growth ratio in the co-culture condition pretreated with L-alanine.

Moreover, co-cultivation with 1% L-alanine can effectively promote the tolerance of lactic acid bacteria to gastrointestinal juice. In the simulated GIT environment, L-alanine has a significant effect on the liability, adhesion and expression of the *LuxS* of the co-cultured bacteria, especially in *L. acidophilus*. while the changes of the auto-aggregation were not significant (Figure 6).

**Figure 6.**
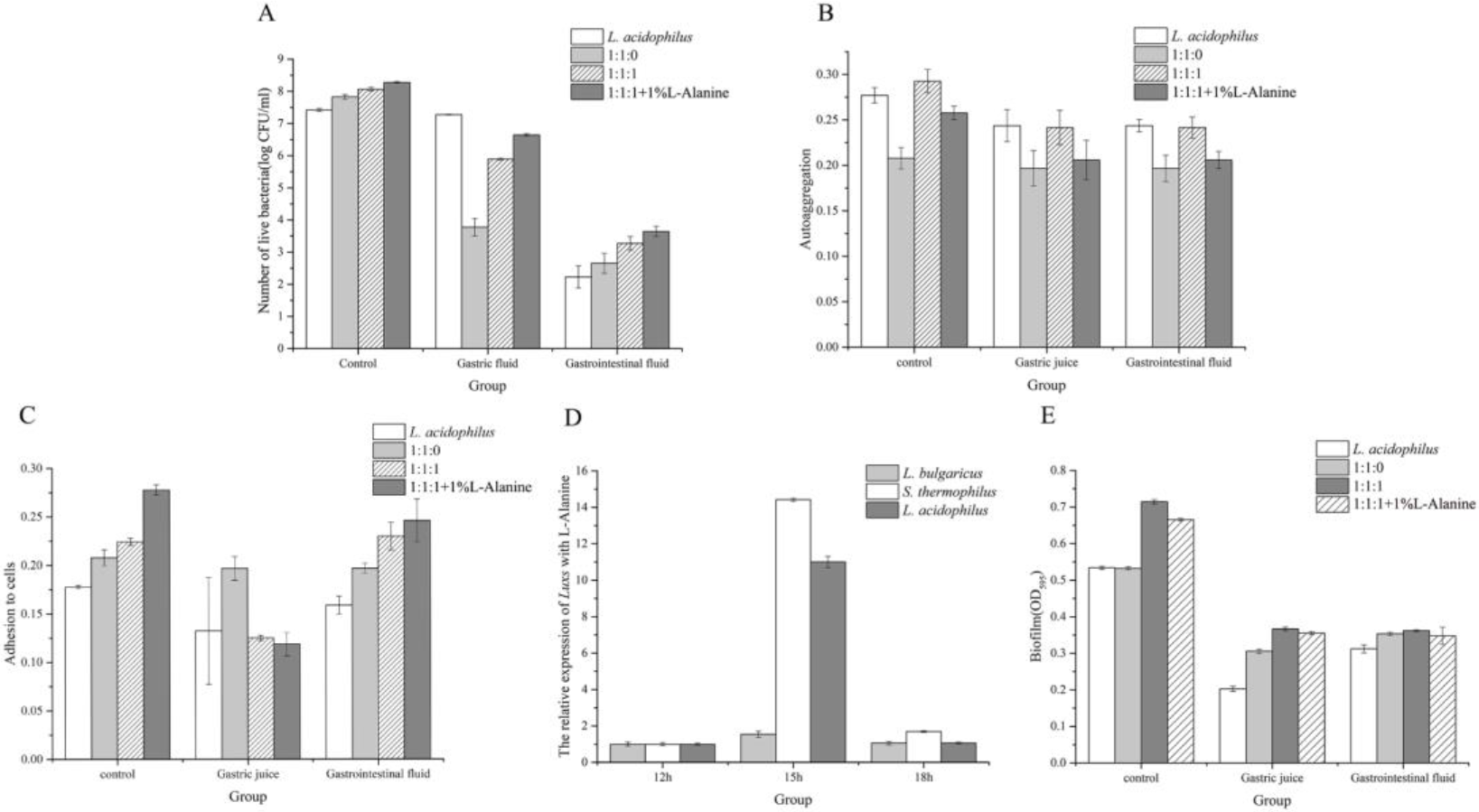
Various physiological indexes of bacteria in the co-culture condition with L-alanine (1%). A, the number of live bacteria in the gastrointestinal fluid under co-culture conditions with 1% L-alanine; B, the autoaggregation rate in the gastrointestinal fluid under co-culture conditions with 1% L-alanine; C, the adhesion to cells in the gastrointestinal fluid under co-culture conditions with 1% L-alanine; D, the relative expression of *LuxS* under co-culture conditions with 1% L-alanine; E, the biofilm formation in the gastrointestinal fluid under co-culture conditions with 1% L-alanine.

### Biofilm formation and liability in the co-culture condition

The results show that biofilm of *L. acidophilus* was significantly reduced after gastrointestinal fluid culture and the co-culture condition can promote the formation of bacterial biofilms. It will also be affected by gastric juice and intestinal juice, but the co-culture condition can make the gastric juice and intestinal juice less affective against the biofilm (Figure 6E). The Live/Dead^®^ BacLight™ cell viability test is an effective method for the detection of the liability and biofilm changes of the bacteria in different conditions, as shown in Figure 7, LAB in the co-culture condition can enhance the cell liability than the mono-culture condition, which is the number of alive bacteria in the 1:1:1 group after stimulated GIT environment treatment were much higher. Meanwhile, when the group co-cultured with 1% L-alanine, the fluorescence intensity was also slightly enhanced.

**Figure 7.**
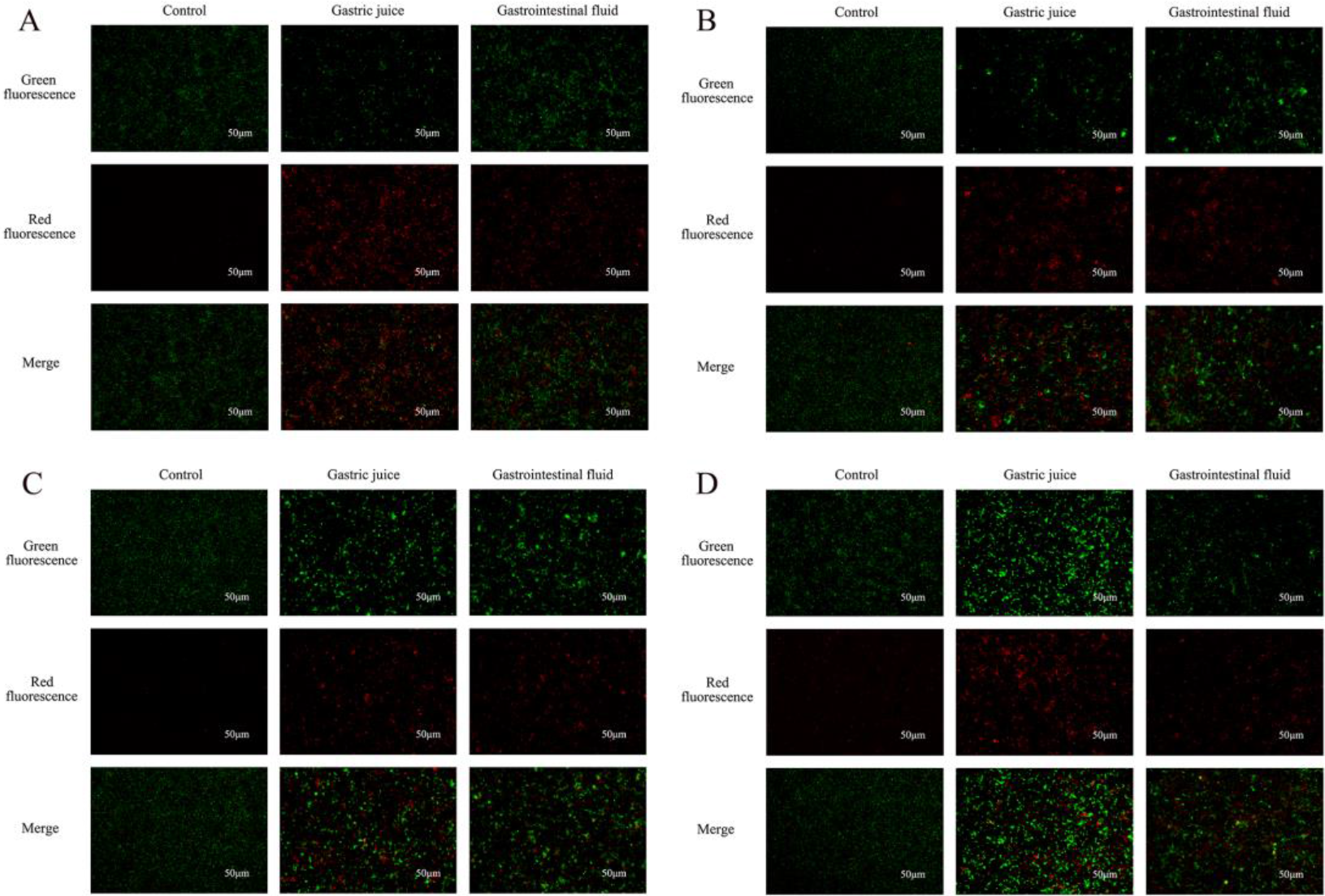
Viability of the bacteria represent with in situ fluorescent biofilm staining. A, the survival of bacteria in *L. acidophilus*; B, the survival of bacteria in 1:1:0; C, the survival of bacteria in 1:1:1; D, the survival of bacteria in 1:1:1 with 1% L-alanine.

## Discussion

As the importance of the probiotics on the beneficial modulation effect of the gut microbiota and its interactions with its host has soared, the use of functional foods containing probiotic bacteria for the health promotion purpose has increased significantly [27]. The most widely used probiotics include LAB, especially *Lactobacillus* and *Bifidobacterium*. While a considerable research focused on the ability of various probiotic strains to beneficially influence host immune responses, metabolic processes and neuro-endocrine pathways [28], the interaction of various probiotic strains with the gut microbes and the host intestinal cells still needs to be studied.

In our study, when *L. bulgaricus*, *S. thermophilus* and *L. acidophilus* were co-cultured at the ratio of 1:1:1, the bacterial density was increased significantly, and the pH value of the co-cultured media also more suitable for the growth of the lactobacillus strains, especially *L. acidophilus*. Previous studies have found that when *L. acidophilus* is co-cultured with *Bifidobacteria*, the environment becomes acidic due to its acid-producing ability which promotes the growth of *L. acidophilus*. At the same time, the co-cultured conditions enhanced the activity of the strain [29]. There is a special way of communication between strains under co-cultured conditions. Quorum sensing is a way of signal transmission between bacteria [30]. In this study, the signal molecules produced were much higher than when they were cultured separately and the participation of *L. acidophilus* also promotes the generation of signal molecules. *L. acidophilus* was reported to promote its intestinal adhesion ability and adaptation by AI-2 quorum sensing [31]. Furthermore, in an experiment of co-cultivating *L. plantarum* DC400 with *L. sanfranciscensis* DPPMA174 or *L. rossiae* A7, the fluorescence intensity of *L. plantarum* DC400 co-cultured with *L. sanfranciscensis* DPPMA174 or *L. rossiae* A7 was significantly higher than that of single culture. At this time, the expression of Luxs gene was 2.5 and 3.5 times that of single culture. This is consistent with our experimental conclusions [32]. During the growth and reproduction of bacteria, signal molecules encoded by specific genes are released. The AI-2 produced by the LuxS enzyme catalyzed by *LuxS* gene encoding between Gram-positive bacteria carries out signal transmission [33]. Our result also proved that during the culture process, the content of *LuxS* gene in *L. acidophilus* was much higher than that of *L. bulgaricus* and *S. thermophilus*. Meanwhile, though the co-culture condition did not promote bacterial proliferation, the co-culture condition can promote the growth of *L. acidophilus*. This is also the reason why the *LuxS* gene are higher than other groups. Due to the limitation of growth space and nutrients *in vitro* the growth curves measured in the flask by the experimental groups are not much different, but the growth of each strain in the co-culture condition was quite different.

Bacteria are known to release a large variety of small molecules, including siderophores, secondary metabolites, and metabolic end products [34]. There are many metabolites containing amino groups in organisms, including protein amino acids, non-protein amino acids, modified amino acids and so on. These amino metabolites cover multiple metabolic pathways in the organism, and they all play important biological roles. For example, in the methionine metabolic pathway, methionine and ATP are activated by methionine adenosyl transferase. Methionine derivatives such as S-adenosylmethionine (SAM), and SAM is the methyl donor for the synthesis of AI-2 [35]. In the metabolite analysis performed using UPLC-MS/MS, L-alanine is screened as the significant amino acid-derived quorum sensing molecule through multivariate statistical analysis and the variable importance of projection. In the KEGG metabolic pathway analysis, L-alanine promoted the synthesis of SAM and S-adenosylhomocysteine (SAH). The addition of 1% L-alanine made the growth of LAB more stable and promoted the growth of *L. acidophilus* and enhanced the expression of the *LuxS* gene level when cultured for 15h in the co-cultured fermentation model.

In addition to the information exchange between LAB, another important feature of LAB strain is their ability to survive gastrointestinal transport [36]. This is related to whether lactic acid bacteria can play a role in the intestine. Fuller (1989) stated that probiotics must be viable microorganisms and produce beneficial effects on their hosts. The results show that the co-culture condition effectively protects the lactic acid bacteria in the gastrointestinal environment. L-alanine can work both as the QS regulator in the synthesis of AI-2 and the enhancer which can increase the GIT tolerance of the lactobacillus strains in the co-cultured fermentation model. Furthermore, in the co-culture condition, the viability, self-aggregation rate, adhesion to cells and biofilm formation of the bacteria were also have a relationship with each other during this special situation. In previous studies, it was found that co-cultured of *L. plantarum* and *P. acidilactici* VTCC 10800 or *S. cerevisiae* Y11-43 significantly improved the biofilm formation ability [38, 39]. In addition, SEM results of biofilms under monoculture and co-culture conditions also confirmed this conclusion. The formation of biofilm can increase the liability of the cell, and the co-culture condition of LAB will provide a for the communication of the bacteria to interact with each other.

## Conclusions

In summary, it is shown that the co-cultured condition is an effective approach for the interaction of the LAB in the fermentation model in this research. Meanwhile, the special and natural fermentation process can promote the release of amino acid-derived quorum sensing molecules which can effectively protect the LAB in the stimulated gastrointestinal environment through the enhancement of the adhesion and biofilm formation. The findings of this study confirmed the QS of the LAB in the co-culture condition *in vitro*, and provide a clue for using the amino acid-derived quorum sensing molecule in the complex fermented dairy industry.

## Acknowledgement

This work was supported by the National Natural Science Foundation of China (32072192, 31671869, 31901668), Key Research and Development Project of Zhejiang Province (2020C02042), the Natural Science Foundation of Ningbo (202003N4129), the Open Project Program of the First-Class Bioengineering Disciplines in Zhejiang Province (KF2020007), State Key Laboratory of Dairy Biotechnology (SKLDB2020-007), the Graduate General Program of the Education Department in Zhejiang Province (Y202045625) and the K. C. Wong Magna Fund in Ningbo University.

